# Blood-based untargeted metabolomics in Relapsing-Remitting Multiple Sclerosis revealed the testable therapeutic target

**DOI:** 10.1101/2021.09.13.460085

**Authors:** Insha Zahoor, Hamid Suhail, Indrani Datta, Mohammad Ejaz Ahmed, Laila M Poisson, Jeffrey Waters, Rui Bin, Jaspreet Singh, Mirela Cerghet, Ashok Kumar, Md Nasrul Hoda, Ramandeep Rattan, Ashutosh K Mangalam, Shailendra Giri

## Abstract

Metabolic aberrations impact the pathogenesis of multiple sclerosis (MS) and possibly can provide clues for new treatment strategies. Using untargeted metabolomics, we measured serum metabolites from 35 relapsing-remitting patients and 14 healthy age-matched controls. Out of 632 known metabolites detected, 60 were significantly altered in relapsing-remitting MS (RRMS). Bioinformatics analysis identified an altered “metabotype” in RRMS patients, represented by 4 changed metabolic pathways of glycerophospholipid, citrate cycle, sphingolipid, and pyruvate metabolism. Interestingly, the common upstream metabolic pathway feeding these 4 pathways is the glycolysis pathway. Real-time bioenergetic analysis of the patient derived peripheral blood mononuclear cells, showed enhanced glycolysis, supporting the altered metabolic state of immune cells. Experimental autoimmune encephalomyelitis mice treated with the glycolytic inhibitor, 2-deoxy-D-glucose ameliorated the disease progression and inhibited the disease pathology significantly by promoting the anti-inflammatory phenotype of monocytes/macrophage in the central nervous system. Our study suggests that targeting glycolysis offers a potential target for MS.

## INTRODUCTION

Multiple sclerosis (MS) is a neuroinflammatory and demyelinating central nervous system (CNS) disease mainly affecting young adults. It is orchestrated by a wide continuum of immune components, which degenerate the axonal myelin sheath in the CNS (Compston and Coles, 2008). The most common disease course includes periods of relapse followed by episodes of partial or complete recovery (remission), known as relapsing-remitting MS (RRMS). Relapses involve acute autoimmune attacks on the brain and spinal cord, leading to multiple demyelinated lesions associated with the infiltration of mononuclear (MN) cells, axonal loss, and neurodegeneration (Compston and Coles, 2008). Despite the tremendous upsurge in the research and development of immunomodulatory drugs for MS in the past two decades, there are no satisfactory treatments that can provide long-term recovery in patients (Torkildsen et al., 2016). The biggest challenge with most of these drugs is substantial side effects and poor tolerance (Saidha et al., 2012). Thus, there is an urgent need to identify novel molecular targets and harness the immune pathways to translate into MS therapies.

Metabolomics is one of the ‘omics’ approaches that uses sophisticated analytical and statistical methods to measure the endogenously produced end products (metabolites) of all the cellular processes in the biological matrices (Zhang et al., 2013). Metabolic profiling can provide a direct window to the instantaneous physiological or pathological changes occurring in a living organism. The art of metabolomics can be implicated to differentiate between a diseased *vs*. a non-diseased state; therefore, it has gained significant translational importance in MS studies for the possible development of MS associated surrogate biomarkers and to monitor the disease progression with or without a therapeutic intervention. A few studies have employed the untargeted metabolomics approach to identify MS patients’ cardinal metabolic changes (Zahoor et al., 2021). However, most of the studies used a single platform with the ability to identify limited metabolites (Zahoor et al., 2021). Previously, we have also performed a comprehensive analysis of plasma and urine metabolites in the experimental autoimmune encephalomyelitis (EAE), wherein we observed alterations in several metabolic pathways and reiterated the role of metabolism in the immune response of MS (Mangalam et al., 2013; Poisson et al., 2015; Singh et al., 2019). Metabolic reprogramming of the immune cells has been reported in the pathogenesis of immune-mediated diseases including systemic lupus erythematosus (Yin et al., 2016), autoimmune arthritis (Weyand et al., 2017), and Guillain-Barré syndrome (Liu et al., 2018). Metabolic reprogramming within the inflammatory cells, such as activated macrophages and CD4 cells, is well documented, where a switch to glycolysis for energy generation is essentially required for their cellular functions (Michalek et al., 2011). The metabolic adaptations and metabolic diversity exhibited by the immune cells provides clues to their selective regulation, by exploiting their specific metabolic requirements (Bettencourt and Powell, 2017). Given that MS is an immune-mediated disease, governed by inflammation, it provides a perfect model for immune cells to undergo metabolic reprogramming.

The purpose of the present study was to characterize and identify the perturbed metabolic pathways in serum samples of RRMS patients and target them using pharmacological approaches to examine their therapeutic potential in disease progression using classical preclinical mouse models of MS.

## RESULTS

### RRMS patients show a distinct serum metabolic profile compared to healthy subjects (HS)

To identify the metabolic signature associated with the disease state in RRMS, we performed untargeted global metabolic profiling in serum collected from 33 RRMS patients and 14 age- and gender-matched healthy subjects (HS). Clinical characteristics of HS and untreated RRMS patients are summarized in **Table 1**. Out of 632 known metabolites measured in RRMS, 60 metabolites mapped to various metabolic pathways were observed to be significantly altered between HS and RRMS (p < 0.05, false discover rate < 0.10), as revealed by Welch’s two-sample t-test (**Table S1**). Among the altered metabolites, 53 (88.33%) metabolites were upregulated, and only 7 (11.67%) downregulated relative to HS. Three-dimensional partial least-squares discriminant analysis of the metabolomics data was performed to further substantiate the observed difference in serum metabolites between HS and RRMS patients. It revealed a clear separation between the 2 groups based on the altered metabolites (**Figure 1A**). To visualize the overall differences between the 60 altered metabolites, a heat map was drawn, which showed a clear difference of metabolite levels between the 2 groups (**Figure 1B**). The pie chart indicates that the most altered metabolites belonged to the lipid (46%) and xenobiotics (20%) pathways, followed by peptide (12%), amino acid (7%), cofactor and vitamin (5%), carbohydrate (5%), energy (3%), and nucleotide (2%) metabolic pathways (**Figure 1C**). Altogether, this data clearly shows that RRMS patients and HS have distinct serum metabolite profiles indicating unique altered metabolic pathways, which can be used to distinguish between RRMS patients and HS.

**Figure 1.**
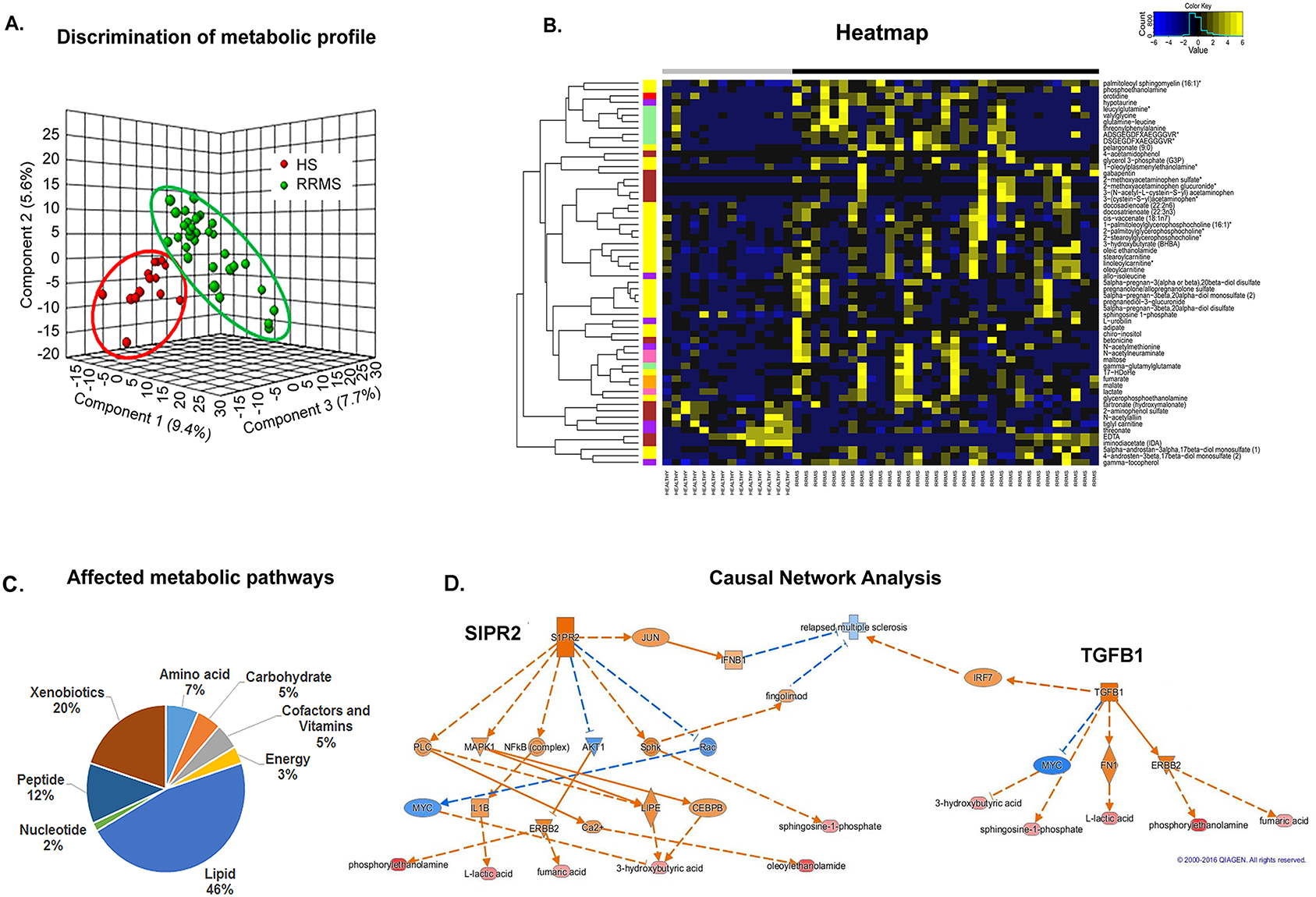
Relapsing remitting multiple sclerosis (RRMS) patients show an altered metabolic state “metabotype” compared to healthy subjects (HS). (A) Three dimensional partial least-squares discriminant analysis (PLS-DA) 3D plots showing significant discrimination of RRMS and HS. (B) Pie chart depicting the classification of metabolic perturbations in the serum of RRMS patients compared with HS. (C) Heat map representative of the hierarchal clustering of the 60 metabolites from each of the replicates of serum from RRMS and HS. Shades of yellow represent elevation of a metabolite and shades of blue represent decrease of a metabolite relative to its mean level in these samples (see color scale). (D) Ingenuity’s IPA software CNA report identified G-protein-coupled sphingosine-1-phosphate receptor 2 (S1PR2) and transforming growth factor-beta-1 (TGFb1) as master regulators predicted to be activated based on the altered levels of metabolites (bottom layer). Shades of orange shows prediction of activation and shades of blue shows prediction of inhibition. Bottom layer of these CNAs are metabolites which are differential between RRMS and HS, while shades of red shows up-regulation in RRMS. Abbreviations:

**Table 1:** Demographic information of healthy and RRMS patients

|  | **Healthy**<br>**(N=14)** | **RRMS**<br>**(N=35)** |
| --- | --- | --- |
| Female, N (%) | 64% | 64% |
| Mean age, years | 40 | 45 |
| Mean disease duration | -- | 18.2 yrs |
| DMTs treatment | NA | None |

DMT, Disease modifying therapies; NA, Not applicable; RRMS, Relapsing-remitting multiple sclerosis.

Ingenuity’s software causal network analysis (CNA; Qiagen; Ann Arbor, MI) (Kramer et al., 2014) was used to identify the underlying signaling cascade of the dysregulated metabolites in RRMS patients and the master regulators of these signaling pathways. CNA showed G-protein-coupled sphingosine-1-phosphate receptor 2 (S1PR2 or S1P2) as one of the master regulators predicted to be activated (S1PR2 activation zscore = 2.449) during disease (**Figure 1D; Table S1**). S1PR2 emerged as the master regulator of a small causal network exhibiting 2 layers of 6 intermediate regulators, which may explain the altered 6 metabolites in RRMS namely phosphorylethanolamine, lactic acid, fumaric acid, 3-hydroxybutyric acid, oleoylethanolamide and sphingosine-1-phosphate shown in the bottom row of the **Figure 1D**. CNA also predicted that RRMS could be repressed indirectly by targeting S1PR2 with JUN, interferon-beta (IFNβ1), and fingolimod as intermediate regulators. Since IFNβ and fingolimod are US Food and Drug Administration approved drugs for MS patients to control disease progression, this further supports the role of S1PR2 and altered metabolic pathways in MS. Another predicted master regulator identified was transforming growth factor-beta-1 (TGFβ1; activation zscore = 2.236). TGFβ1 was connected to regulating a small causal network of 3 intermediate regulators; Myc, fibronectin 1 and erythroblastic oncogene B, resulting in an alteration of 5 metabolites in RRMS (**Figure 1D**). TGFβ contribute to the MS pathogenesis in combination with another gene: interferon regulatory factor 7 (IRF7) (**Figure 1D**). To gain further insight into the signaling nodes and pathways connected with the altered metabolites, we also performed Ingenuity Pathway Analysis (IPA). To be noted, the top canonical pathway in IPA was also sphingosine and sphingosine-1-phosphate metabolism (p < 4.34E-03; **Figure S1**). Further the top network (Cell signaling, molecular transport, vitamin and mineral metabolism; score 33 with 13 focus molecules; **Figure S2**) also revealed a signaling hub involving SIPRs. Thus, all 3 analyses revealed an association with the common metabolic pathway of sphingosine-1-phosphate metabolism, which is known to play a key role in regulating inflammation in MS (Cohan et al., 2020). Thus, our metabolic profiling of RRMS patients corroborates and validates that metabolite alterations are reflective of the biological processes underlying the disease.

Further, to identify and visualize the enriched metabolic pathways from dysregulated metabolites in RRMS, we employed Metscape, a plug-in for Cytoscape (Gao et al., 2010). Metscape extracts and integrates metabolite database records from several public sources and provides a complete view of entire metabolic networks, including the connection between metabolites and genes, compounds networks, and also provides information for reactions, enzymes, and associated pathways. A number of pathway networks were enriched in RRMS patients compared to HS (**Figure 2A**). Metabolic networks including glycolysis, tricarboxylic acid cycle (TCA), urea, purine, tyrosine, butnoate, phosphatidylinositol phosphate, glycerosphingolipids, pyrimidine, aminosugar, methionine and cysteine were observed to be greatly impacted during MS disease and are reflected in blood (**Figure 2A**). To further understand the functional role of metabolic alterations in RRMS and their link to various metabolic pathways, the Kyoto Encyclopedia of Genes and Genomes (KEGG) metabolic library was analyzed using MetaboAnalyst (**Figure 2B**) as previously described (Mangalam et al., 2013; Poisson et al., 2015; Singh et al., 2019). Results of each of the 80 human pathways of KEGG were simultaneously plotted to show the most significant pathways in terms of Global Test p-value (vertical axis, shades of red) and impact value (horizontal axis, circle diameter). The top 4 pathways found to be linked with altered metabolites by p-value or impact included 1) glycerophospholipid metabolism, 2) citrate cycle (TCA), 3) sphingolipid metabolism, and 4) pyruvate metabolism (**Figure 2B-C**). The glycerophospholipids are glycerol-based phospholipids, and their primary function is to serve as a structural component of the biological membrane. On the other hand, sphingolipids protect the cell surface against harmful environmental factors by forming a mechanical and chemically resistant outer leaflet of the plasma membrane lipid bilayer. Sphingolipid metabolites such as S1P and ceramides have been known to play an important role in various functions under normal or pathological conditions (Maceyka and Spiegel, 2014). Citrate cycle, also known as TCA or Krebs cycle, is an important aerobic pathway to generate energy, as cell needs to grow and divide. Pyruvate metabolism is a key player in balancing energy metabolism in the body.

**Figure 2.**
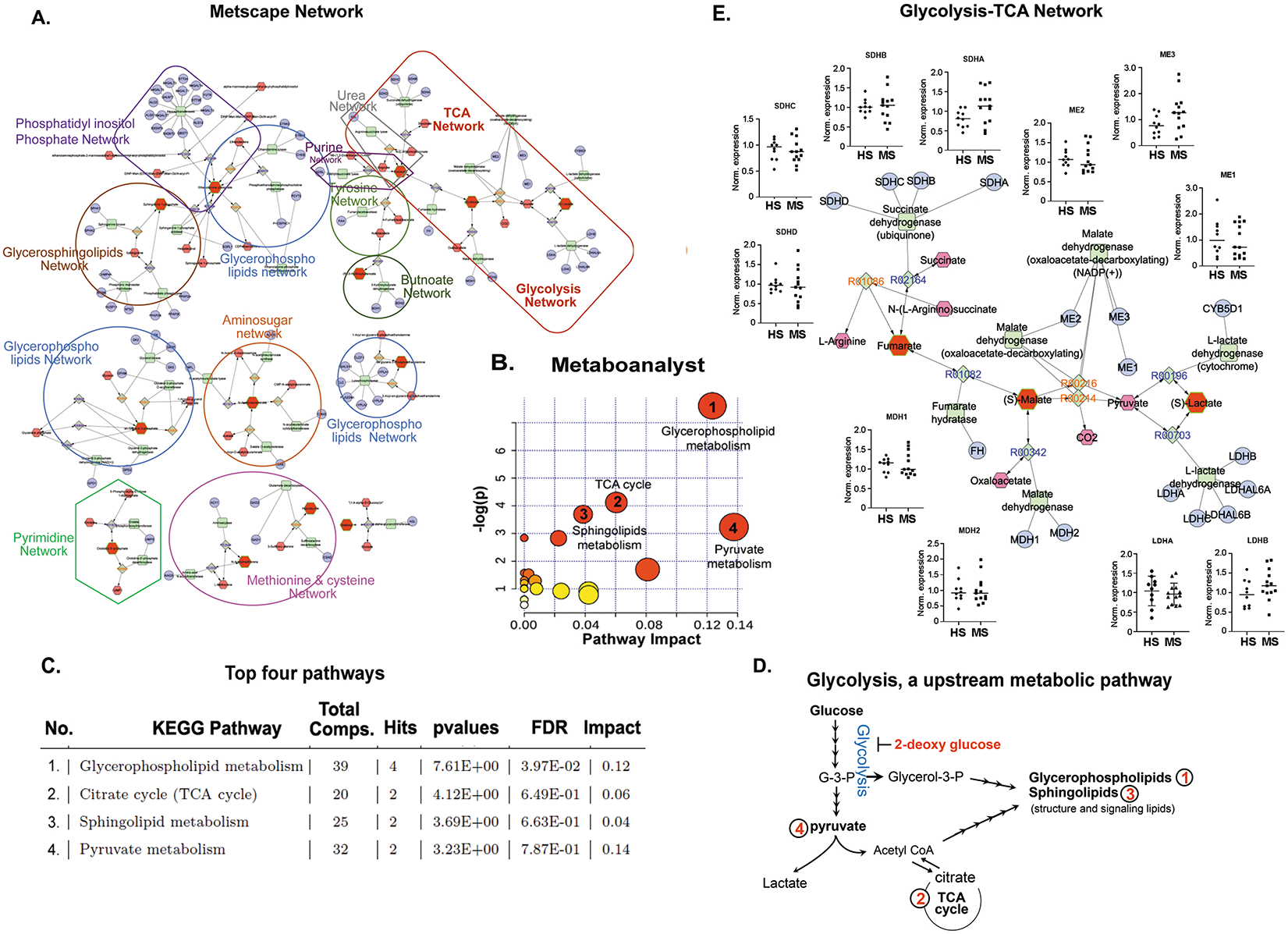
Bioinformatics analysis identified glycolysis as an upstream to the altered metabolic pathways in RRMS. (A) Visualization and interpretation of metabolomic network of RRMS patients using Metscape in the context of human metabolic networks. In Metscape, compounds are represented as hexagons, reactions are diamonds, enzymes are squares, and genes are circles. Input metabolites are represented as red hexagons with green border. (B) Sixty metabolites that displayed significantly different abundance between RRMS and HS were subjected for the pathway enrichment analysis (y axis, enrichment p values) and the pathway topology analysis (x axis, pathway impact values, and indicative of the centrality and enrichment of a pathway) in the pathway module of MetaboAnalyst 4.0. The color of a circle is indicative of the level of enrichment significance, with red being high and yellow being low. Bigger size of a circle is proportional to the higher impact value of the pathway. (C) The top 4 pathways that arise with low p-values and with high impact are indicated in the table format. (D) Schematic indicating glycolysis is upstream to the 4 altered metabolic pathways identified in RRMS. (E) Schematic depicting the comparative analysis of gene expression between HS and RRMS cases for the enzymes involved in glycolytic-TCA pathway, linked with altered metabolites found in RRMS.

Based on the current biochemistry knowledge, all 4 identified dysregulated pathways are interconnected and point towards glycolysis as an upstream metabolic pathway (**Figure 2D-E**). Glycolysis is a sequence of 10 chemical reactions taking place in most of the cells that break down glucose, with the production of pyruvate or lactic acid and releasing energy that is captured and stored as adenosine triphosphate (ATP). Glycolysis is the major metabolic pathway that along with energy provides metabolites and mediators for various biosynthesis pathways that are essential for providing biomass for a healthy proliferating and functional cell. Pyruvate is a key player in energy metabolism and could be converted into lactate by lactate dehydrogenase (LDH) and released to extracellular space, or it could be converted to acetyl-CoA by pyruvate dehydrogenase and enter the TCA cycle, which could serve as a precursor for the synthesis of lipids (**Figure 2D**). Another intermediate of glycolysis, dihydroxyacetone phosphate, gives rise to glycerol 3-phosphate, a reaction catalyzed by glycerol-3-phosphate dehydrogenase. Glycerol 3-phosphate is essential for *de novo* synthesis of glycerophospholipids, sphingolipids, and lysolipids (**Figure 2D**). Thus, glycolysis is the upstream feeder metabolic pathway of all the 4 perturbed metabolic pathways identified in RRMS patients.

Since glycolysis-TCA network emerged as the prominent altered metabolic pathway (**Figure 2E**), we examined the messenger RNA expression of the key enzymes involved in the reactions of the differential altered metabolites in the peripheral blood mononuclear cells (PBMCs) of healthy and RRMS patients. Demographic detail of this cohort is listed under Table 2. We did not observe any change in the expression of lactate dehydrogenase A and B (LDHA and LDHB), malate dehydrogenase-1 and −2, melic enzyme-1, −2 and −3 and succinate dehydrogenase-A, -B, - C and -D enzymes (**Figure 2E**). To further corroborate our findings, we used 3 datasets from the National Center for Biotechnology Information (NCBI) Gene Expression Omnibus (GEO) database, including GSE21942 (Kemppinen et al., 2011), GSE26484 (Nakatsuji et al., 2012), and GSE43591 (Jernas et al., 2013), and the raw data of each dataset was preprocessed with the Robust Multiarray Average (RMA) algorithm. Using Limma (or geo2R) differential expression analysis was performed on each dataset. Genes with an adjusted P-value lower than 0.05 considered as significant. We analyzed the expression of genes involved in glycolysis-TCA pathway (**Figure 2E**) and found that none of these genes were affected in MS patients compared to HS; except the LDHA and ASL genes in GSE21942 (**Table S2**) (Kemppinen et al., 2011). Overall, our data suggests that glycolysis-TCA pathways are being altered in the PBMCs of MS patients without affecting their gene expression, indicating a regulation of their function or activity by the inflammatory environment of the disease.

**Table 2:** Demographic information of healthy and RRMS patients used for qualitative polymerase chain reaction data for metabolic genes presented in figure 2D.

|  | **Healthy**<br>**(N=10)** | **RRMS**<br>**(N=13)** |
| --- | --- | --- |
| Female, N (%) | 60% | 77% |
| Mean age, years | 43 | 44.9 |
| Mean disease duration, years | -- | 4.5 |
| DMTs | NA | None |

DMTs, Disease modifying treatments; NA, not applicable; RRMS, Relapsing-remitting MS.

### PBMCs from RRMS patients have increased glycolysis compared to HS

Modulation of the glycolysis pathway is reported to be directly related to the inflammatory response in immune cells (O’Neill and Pearce, 2016). Our data also suggests an altered glycolysis metabolism in the PBMCs of MS patients, which can be postulated to result in an altered immune response and neuroinflammation. We performed the glycolytic response profile, measured as extracellular acidification rate (ECAR), in HS and RRMS patient derived PBMCs using the Seahorse XF^e^96 analyzer (Agilent, Santa Clara, CA). PBMCs from both HS and RRMS responded to the various stressors and gave a typical glycolytic profile (**Figure 3A**). However, the basal glycolysis was significantly (p < 0.05) higher in the RRMS PBMCs when compared to HS, indicating an inherent increase in the glycolytic activity of the RRMS PBMCs (**Figure 3A**). This observation was also replicated in the preclinical mouse model of EAE, where PBMCs of the EAE group exhibited higher levels of glycolysis compared to control complete Freund’s adjuvant (CFA) groups (**Figure 3B**). These findings explicitly show that the PBMCs from both MS patients and from its preclinical mouse perform glycolysis at an increased rate.

**Figure 3.**
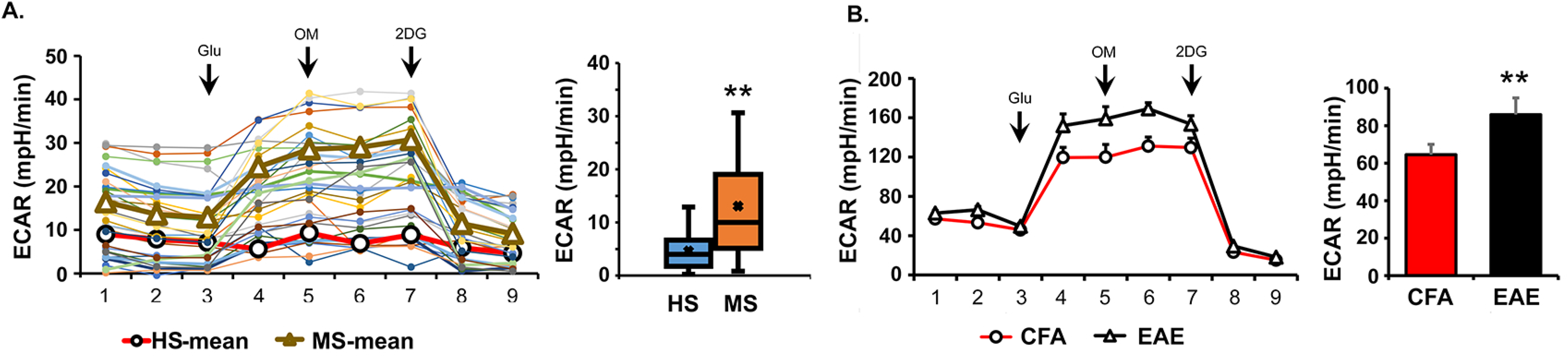
Higher glycolytic activity was observed in the peripheral blood mononuclear cells (PBMC) of RRMS patients and in preclinical mouse model (EAE). (A) Isolated PBMC of RRMS patients (N = 17) and HS (N = 14) were analyzed for glycolysis (ECAR) measurement using Seahorse Bioanalyzer. (B) PBMC of EAE and CFA groups were analyzed for glycolysis measurement using Seahorse Bioanalyzer (mean + SEM, N = 6). **p < 0.01; **p < 0.05 compared to HS. Abbreviations:

### Targeting glycolysis using 2-deoxy-glucose treatment ameliorated disease progression in preclinical EAE mouse models

Since glycolysis is upregulated in PBMCs and emerged as the upstream metabolic pathway regulating/connecting/feeding into the 4 altered metabolic pathways identified in RRMS patients, we questioned if inhibiting glycolysis will impact MS disease progression. We employed 2-deoxy-glucose (2DG), a non-metabolizing glucose analog and a competitive inhibitor of hexokinase, to examine the effect of glycolysis inhibition in the relapsing-remitting preclinical model of MS using SJL mice. Mice immunized for EAE with proteolipid protein (PLP) were treated daily with 2DG (50 mg/kg, intraperitoneally) from day 6 post-immunization, while the control group was injected with phosphate buffered saline (PBS). Treatment with 2DG significantly reduced the severity of the disease (2.7 ± 0.22 vs 1.2 ± 0.54; at peak of the disease, p < 0.01) without affecting the onset of the disease (**Figure 4A)**. Treatment with 2DG also abrogated the relapse in the treated group compared to the PBS group (1.9 ± 0.11 vs 0.2 ± 0.22 at day 25 post-immunization; p < 0.001, (**Figure 4A**). Similar efficacy of 2DG was observed in 2 other mouse models of EAE, where 2DG was effective in not only in delaying the disease onset but also reduced severity (2.6 ± 0.21 vs 0.3 ± 0.14; p < 0.001 in B6 model and 2.4 ± 0.27 vs 0.3 ± 0.22; p < 0.001 in 2D2 TCR transgenic model) (**Figure 4B-C**). We also examined the effect of 2DG when given orally in the 2D2 EAE model via drinking water at the dose of 50 mg/kg (1.25 mg per 6 ml as a mouse drink 6-7 ml per day) from day 6 post-immunization. The treated group with 2DG showed delayed disease onset and reduced disease severity compared to the untreated EAE group (**Figure 4D**), demonstrating the positive impact of 2DG in reducing the disease progression in EAE mouse models. A decrease in clinical score of treated EAE was reflected by the reduction in the number of infiltrated MN cells and preservation of myelin content as examined by the hematoxylin and eosin and Luxol fast blue staining, respectively, in the spinal cord sections (**Figure 4E-F**). As evident from the hispothaological analyses, there is significant reduction in the infiltration of immune cells and lesion size in 2DG treated mice as compared to vehicle treated EAE mice. Moreover, demyelination also was sigificantly prevented in the 2DG treated group compared to vehicle/PBS treated EAE mice. Therefore, our findings strongly advocate that 2DG treatment following EAE reduces immune cell infiltration, prevents lesion progression and promotes remyelination in the preclinical models of MS disease.

**Figure 4.**
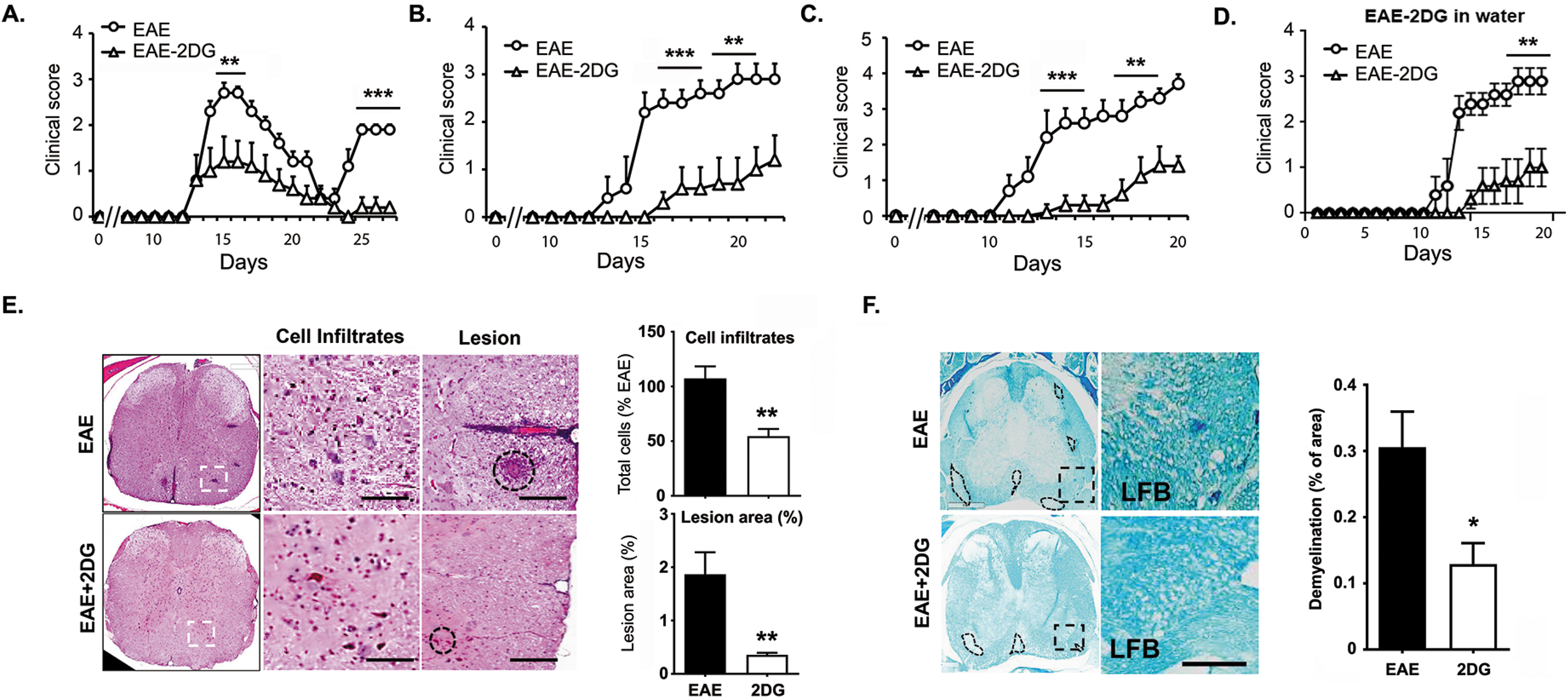
Targeting energy pathway ameliorates disease progression in EAE mouse models. (A-C) EAE was induced in SJL, B6 and 2D2 mice using PLP and MOG 35-55, respectively. One set of the group was given daily 2-DG (50 mg/kg body weight, intraperitoneally in 200 ul volume) and another set was given phosphate buffered saline as vehicle. Clinical score was taken till the end of the study (N = 15). **p < 0.01; ***p < 0.001 compared to vehicle treated EAE group was considered as statistically significant. (D) EAE was induced in 2D2 mice with MOG and one set was given 2DG in drinking water (0.2 mg/ml; w/v) from day 7 post-immunization. Clinical score was taken till the end of the study (N = 6). **p < 0.01 compared to vehicle treated EAE group was considered as statistically significant. (E-F) Representative images shows histopathological changes in the spinal cord tissues in EAE mice with or without 2DG treatment. Sections were staind with hematoxilin and eosin (H&E) to show cells infiltration and lesion size and Luxol fast blue to (LFB) to demonstrate changes in myelin content. Data were represented as mean ± SD (n = 6 mice/group). Scale bar, 100-μm. Statistical analyses were done with Student’s t-test. Statistical significance was determined at P < 0.05. Abbreviations: EAE,; SD, standard deviation.

Further, to examine the effect of 2DG on myelin-specific immune response, we isolated spleen cells from 2DG treated and vehicle-EAE groups at day 20 post-immunization and stimulated them with myelin oligodendrocyte glycoprotein (MOG)_35-55_ (20 μg/ml) for 72 hours. Cell proliferation was examined after 96 hours by cell proliferation assay (Promega, Madison, WI), and cell supernatants were used for enzyme-linked immunosorbent assay to see the status of inflammatory cytokines in both groups. 2DG treatment significantly reduced antigen-specific cell proliferation (data not shown), reduced the levels of IFNγ, interleukin (IL) 17, and granulocyte-macrophage colony-stimulating factor (GM-CSF) and induced the production of IL10 compared to untreated EAE group (**Figure 5A**). Consistent with the protein levels, quantitative polymerase chain reaction (qPCR) data showed that the messenger RNA expression of all cytokines was reduced by 2DG treatment (**Figure 5B**). Infiltration of MN cells, including myeloid cells and T cells that inflame the CNS environment, is detrimental for oligodendrocytes and neurons, leading to demyelination in MS and EAE. To examine the potential mechanism of 2DG-mediated protection in EAE, we quantified the absolute number of infiltrating MN cells in the CNS and their ability to produce pro-inflammatory cytokines. We observed that 2DG treatment significantly reduced the overall number of infiltrating MN cells (CD45^+^) and CD4 T cells (CD45^+^CD3^+^CD4^+^) (**Figure 5C**) as examined by flow cytometry and immunohistochemical staining for CD4 (**Figure 5D**). Along with the reduction of infiltration, the CD4 T cells also showed reduction in the intracellular pro-inflammatory cytokines being produced, including IFNγ, IL17, and GMCSF (**Figure 5E-F**). Thus inhibition of glycolysis by 2DG ameliorates the EAE disease progression in various preclinical models and is able to reduce the number and capacity of the infiltarting immune cells.

**Figure 5.**
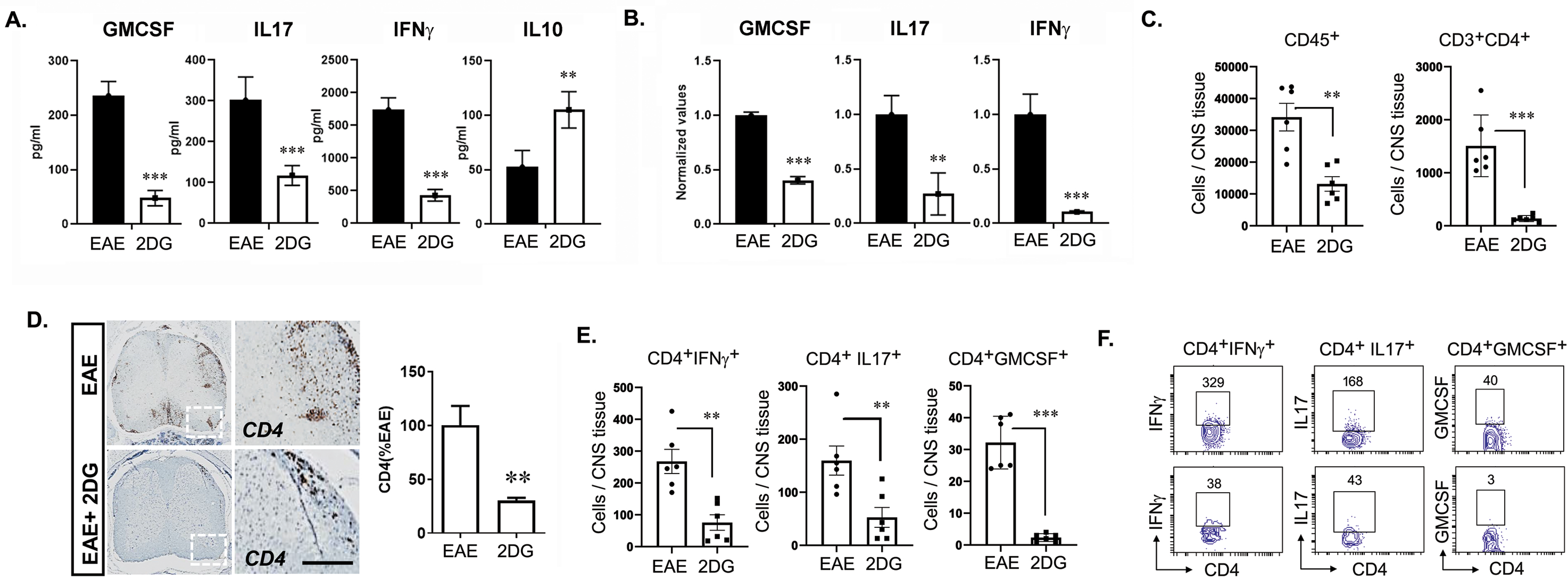
Targeting energy pathway moldulates CD4 cells infiltration and response in EAE mouse. (A) On day 20^th^, splenocytes of EAE treated and untreated with 2DG in drinking water were stimulated with MOG 35-55. Post 72 hours, proinflammatory (GM-CSF, IL17 and IFNɣ) and anti-inflammatory (IL10) wer examined in cell supernatant using their specific enzyme-linked immunosorbent assay. The data presented are the mean ± SD of 2 indpedent exeriments (N = 5 per group). (B) Expression of GM-CSF, IL17 and IFNɣ were examined in splenocytes of both groups using quantitative PCR and normalized with the control gene; ribosomal L27 housekeeping gene. The data presented are the mean ± SD of 4 values. (C) Quantitation of CD45+, CD4+, Ly6C^+^ and F4/80^+^ cell number in the spinal cord and brain of EAE and 2DG treated (drinking water) groups. The data presented are the mean ± SD of 6 animals. (D) Representative immunohistochemical staining and quantitative analysis for CD4^+^ T-cell infiltration in EAE induced mice with or without 2DG treatment in drinking water. Data were shown as mean ±SD (N = 6 mice/ group). Scale bar,100-µm. Statistical analysis were done by student’s t test. Statistical significance was determined at P < 0.05. (E-F) Quantitation of IFNɣ, IL17a and GM-CSF producing CD4+ T cell number in the spinal cord and brain of EAE and 2DG treated (drinking water) groups. The data presented are the mean ± SD of 6 animals. N = 4-6 per group; mean + SEM. *p < 0.05, **p < 0.01, and ***p < 0.001. Student’s t test, one-way analysis of variance. Abbreviations:

### Inhibition of glycolysis by 2DG restored the metabolic programming of monocytes/macrophages in EAE

Monocytes/macrophages are the major effectors of demyelination in both MS and EAE and are highly plastic. Once monocytes infiltrate into CNS during disease, depending upon the local environment, monocytes differentiate either into pro-inflammatory (M1) or anti-inflammatory (M2) type macrophages, and their ratio determines the outcome of EAE disease. We examined the effect of 2DG on the infiltration and phenotype of monocytes/macrophages in CNS at day 20 post-immunization. We observed that 2DG treatment significantly reduced the number of Ly6C^+^ monocytes (CD45^+^Ly6C^+^Ly6G^-^) and macrophages (CD45^+^Ly6G^-^F4/80^+^) in the treated group compared to untreated EAE group (**Figure 6A**). This observation was also supported by the immunohistochemical staining for F4/80 in the lumbar area of the spinal cord sections (**Figure 6B**). 2DG incorporates into the cells and is phosphorylated by hexokinase, resulting in a non-metabolizing 2 deoxy-glucose-6-phosphate (2DG6P), thereby inhibiting glycolysis. To test if activated monocytes take up 2DG during disease, we isolated monocytes from the spleens of 2DG-treated and untreated EAE groups using a monocyte isolation kit (StemCell Technologies, Waltham, MA) on day 20 post-immunization and estimated the levels of 2DG6P. The monocytes from the 2DG treated group had significantly higher levels of 2DG6P (**Figure 6C**), suggesting that in treated animals 2DG is taken up by monocytes. According to our hypothesis this would mean that the monocytes isolated from the treated group would exhibit a metabolically altered phenotype represented by an altered glycolysis. Indeed we observed that monocytes isolated from 2DG treated EAE mice demonstrated reduced glucose uptake (**Figure 6D**) and decreased lactate secretion in media (**Figure 6E**), indicating an inhibition of glycolysis by 2DG treatment.

**Figure 6:**
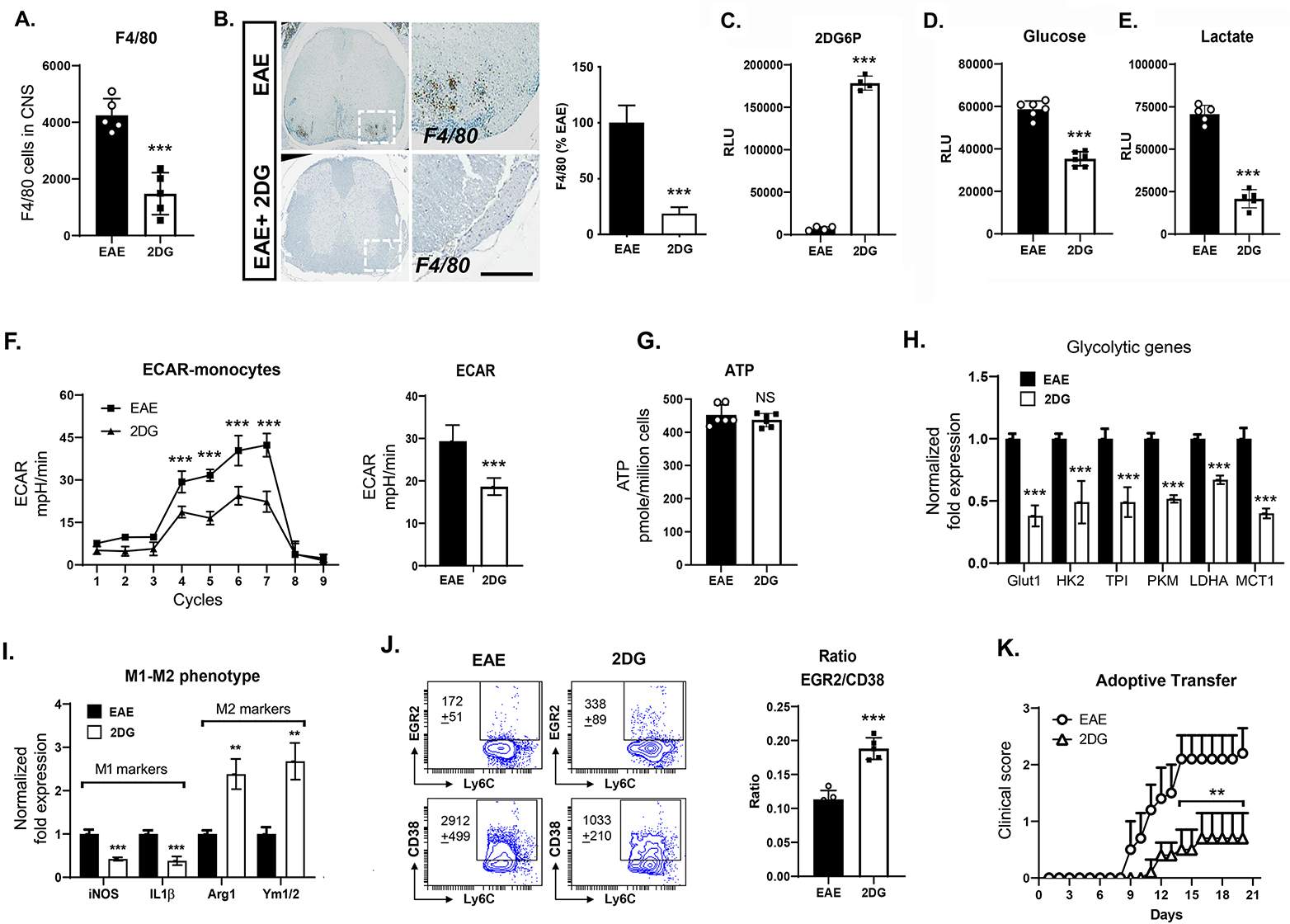
2-Deoxy-glucose induced anti-inflammatory phenotype of monocytes by abrogating glycolytic metabolic pathway. (A-B) Representative immunohistochemical images and quantitation of F4/80^+^ infiltrated macrophage numbers in the spinal cord of EAE mice with or without 2DG treatment. The data were presented as the Mean ±SD (N = 6 mice/group). Scale bar,100-µm. Statistical significance was determined at P < 0.05. (C) Isolated monocytes (20 K) from spleen cells from both groups were processed for the detection of 2 deoxy-glucose-6-phosphate (2DG6P) using Glucose Uptake-Glo^tm^ Assay kit from Promega. The data presented are the mean ± SD of 4 values. (D-E) Isolated monocytes ( > 92%) from spleen cells from both groups were plated (20 K per well) in 96 well plate and post 24h, cell cupernatant was processed for glucose and lactate measurment for determination of glucose consumtion and lactate release in media. The data presented are the mean ± SD of 6 values. (F) Isolated monocytes from spleen cells from both groups were processed for glycolysis measurement using Seahorse Bioanalyzer (N = 8). (G) ATP was measured in isolated monocytes using ATPlite luminescence assay kit from Perkin Elmer. The data presented are the mean ± SD of 6 values. (H) Expression of glycolytic genes including glucose transporter 1 (Glut 1), hexokinase 2 (HK2), triose-phosphate isomerase (TPI), pyruvate kinase M (PKM), lactate dehydrogenase A (LDHA) and monocarboxylate transporter 1 (MCT1) were examined in isolated monocytes from both groups using quantitative PCR. Expression of glycolytic genes were normalized with the control gene; ribosomal L27 housekeeping gene. The data presented are the mean ± SD of 4 values. (I) Expression of M1 [inducible nitric oxide synthase (iNOS) and interleukin 1 beta (IL1β)] and M2 [arginase 1(Arg 1) and chitinase-like 3 (Ym1/2)] specific genes were examined in isolated monocytes from both groups using quantitative PCR and normalized with the control gene; ribosomal L27 housekeeping gene. The data presented are the mean ± SD of 4 values. (J) The ratio of EGR2 and CD38, a marker for M2 and M1 were examined on Ly6C^+^ cells gated on CD11b^+^Ly6G^-^ population by flow cytometry in the spleen of treated and untreated EAE group with 2DG on day 20 post-immunization. (K) Adoptive transfer of monocytes from EAE and 2DG treated EAE groups on day 6 in B6 mice immunized with MOG_35-55_. The data presented are the mean ± SEM of 5 animals. N = 4-6 per group; *p < 0.05, **p < 0.01, and ***p < 0.001. student’s t test, one-way ANOVA. Abbreviations: ATP, adenosine triphosphate

To further determine the effects of 2DG on macrophage metabolism *in vivo*, on day 20 post-immunization splenic macrophages were isolated and processed for glycolysis measurement. We observed that isolated macrophages from untreated EAE exhibited an increased glycolytic rate, which was significantly inhibited upon 2DG treatment *in vivo* (**Figure 5F**). Unchanged total cellular ATP levels suggested that enhanced glycolysis in either group’s macrophages is not contributing to energy production (**Figure 6G**). Increased glycolysis, as a provider of biomass, supports the *de novo* synthesis of fatty acids for the expansion of the endoplasmic reticulum and Golgi, which is required for the production and secretion of proteins integral to APC activation (Everts et al., 2014). Macrophages from vehicle-treated mice showed a ‘higher metabolic phenotype,’ while the 2DG-treated macrophages showed a ‘lower metabolic state,’ similar to that of healthy control macrophages (data not shown). Decreased glycolysis in monocytes/macrophages isolated from 2DG treated EAE was associated with reduced expression of key glycolytic enzymes including glucose transporter 1, hexokinase 2, triose-phosphate isomerase, pyruvate kinase M, LDHA and monocarboxylate transporter 1 compared to the untreated group (**Figure 6H**).

As increased glycolysis is considered a characteristic of M1 macrophages, we observed that splenic macrophages isolated from mice with EAE treated with 2DG displayed an M2 phenotype (higher expression of arginase 1 and Ym1/2) and reduced M1 phenotype (decreased expression of inducible nitric oxide synthase [iNOS] and IL1), resulting in a higher ratio of M2/M1 monocytes/macrophages phenotype (**Figure 6I**). This observation is further supported by flow cytometry where ratio of EGR2 and CD38 is significantly increased in macrophages from 2DG treated group compared to untreated EAE (**Figure 6J**). Anti-inflammatory (M2) macrophages are protective in EAE, and since 2DG polarized macrophages towards the M2 anti-inflammatory phenotype, we next examined if monocytes isolated from 2DG treated mice can modulate EAE disease. To test this, monocytes (1×10^6^) isolated from 2DG-treated or untreated EAE groups were adoptively transferred into recipient B6 mice with active EAE on day 6 post-immunization. The adoptively transferred 2DG-treated monocytes significantly ameliorated the severity of EAE compared to the group of adoptively transferred monocytes from untreated EAE into recipient MOG immunized B6 mice (**Figure 6K**). These data strongly suggest that inhibition of glycolysis by 2DG polarizes monocytes/macrophages towards an anti-inflammatory M2 phenotype, which have the capability to suppress the EAE disease course.

### Inhibition of glycolysis by 2DG restricts neuroinflammation

To further delineate the mechanism of 2DG-mediated protection, we tested if 2DG treatment impacts the MOG-primed MN cell’s ability to induce inflammation in the CNS. We used in vitro co-culture model, where we treated spleen/lymph node single suspension cells with MOG_35-55_ in the presence or absence of 2DG (1 mM) *in vitro*. After 72 hours, conditioned media (CM) and isolated MN cells from both treatment groups (MOG-CM and MOG-2DG-CM) were used for further studies (**Figure 7A**). First, isolated MOG-primed or 2DG treated MOG-primed MNs cells were cocultured with primary brain glial cells (4:1 ratio of MNs cells:mixed glial cells) and post-24-hours, co-cultured cells were assessed for inflammatory markers and mediators. Mixed glial cells co-cultured with MOG-MNs showed increased expression of iNOS and production of nitric oxide (NO), while, 2DG-treated MOG-primed MNs cells were unable to induce iNOS expression in the mixed glial cells (**Figure 7B**). Similarly, MOG-primed MNs cells induced the expression of proinflammatory cytokines including IL1β, TNFα and IL6 in mixed glial cells; however, 2DG treated MOG-primed MNs could not induce cytokines production when cocultured with mixed glial cells (**Figure 7C**). Similar to MOG-primed MNs mixed glial co-culture system, MOG-CM also induced the expression of inflammatory mediator (iNOS) and cytokines (IL1β, IL6 and TNFα) in the mixed brain glia cells while 2DG-CM inhibited the iNOS expression and cytokines production when added in culture media of mixed glia (1:1 ratio) compared to MOG-CM (**Figure 7D**). These sets of data suggest that 2DG altered MN cells could not stimulate proinflammatory cytokines and iNOS expression in mixed glial cells. Further, to examine the effects on oligodendrocytes, rat primary oligodendrocytes were cultured in MOG-CM and 2DG-CM for 24 hours and processed for lactate dehydrogenase activity, a marker for cell death. MOG-CM induced high levels of LDH, which was significantly reduced by 2DG-CM. Cells were processed for MTT assay, which measures cellular metabolic activity as an indicator of cell viability and cytotoxicity. We observed that MOG-CM induced cell death in the oligodendrocytes; however, 2DG-treated MOG-CM was protective (**Figure 7E**). Overall, these data strongly show that inhibition of glycolysis by 2DG alters the MN function and phenotype resulting in protection from inflammation and demyelination.

**Figure 7:**
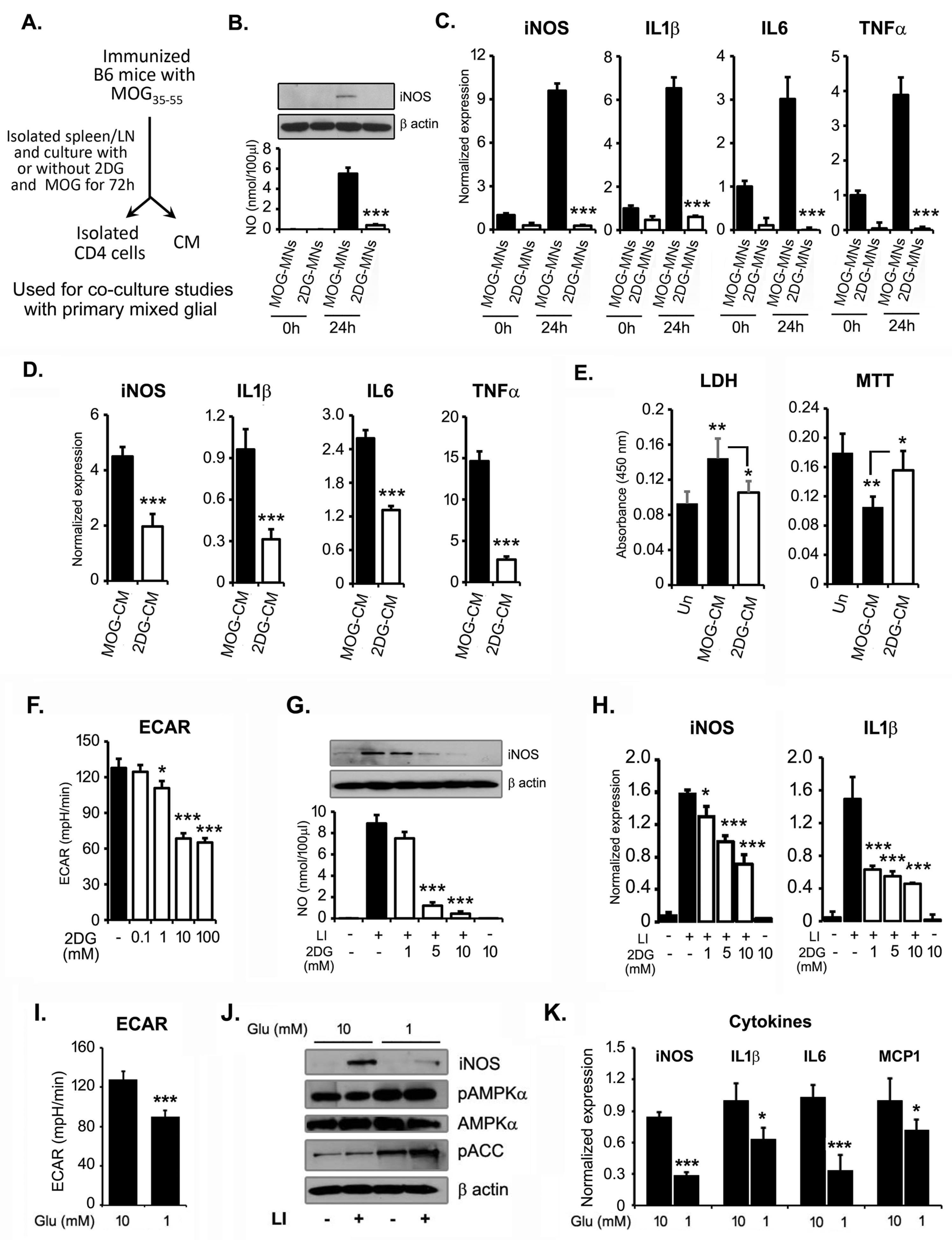
Reducton in glycolysis inhibits activation of brain glial cells. (A) Schematic flow of experimetal design to generate conditioned media (CM) and activated mononuclear cells (MNs) cells in the presence or absence of 2DG (1mM) and used for further experiments. (B-C) For *in vitro* co-culture model, treated and untreated MOG-primed MNs cells with 2DG were cocultured with primary brain glial cells (4:1 ratios of MNs cells: mixed glial cells) for 24h and were processed for nitic oxide (NO), iNOS and cytokines expression by immunoblot and qPCR. The data presented are the mean ± SD of 3 values. (C) MOG-CM and 2DG-CM were added in culture media of mixed glia (1:1 ratio with serum free RPMI) and expression of inflammatory mediator (iNOS) and cytokines (IL1β, IL6 and TNFα) was examined by qPCR. (D) MOG-CM and 2DG-CM were added in the culture media of isolated rat oligodendrocyte precursor cells (OPCs) in the ratio of 1:1 with serm-free DMEM and LDH and MTT assays were performed post 24h of incubation. The data presented are the mean ± SD of 6 values. (F) Dose-dependent effect of 2DG (0.1-100 mM) on ECAR in brain glial cells using Seahorse Bioanalyzer (N = 6-8). (G-H) Brain glial cells were treated with 2DG (1-10 mM) for 30 minutes prior the treatment of LI (0.5µg/10ng per ml). Post 18h, cells were processed for iNOS immunoblot analsis and cell supernatant was processed for NO. Under similar experimental condition, at 6h of LI stimulation, cells wer processed for RNA isolation and qPCR for the detection of iNOS and IL1β. The data presented are the mean ± SD of 3 values. (I) Impact of glucose on ECAR in brain glial cells using Seahorse Bioanalyzer (N = 6). (J) Primary brain glial cells were cultured in normal (10 mM) and low (1 mM) glucose condition in RPMI for 24h followed by stimulation with LI for 18h. The levels of iNOS, pAMPKα, AMPKα and pACC were examined by immunoblot analysis using their specific antibodies. Beta actin was used to examine the equal protein loading. Represented blots are the one of the 2 independent experiments. (K) Primary brain glial cells were cultured in normal (10 mM) and low (1 mM) glucose condition in RPMI for 24h followed by stimulation with LI for 6h. Cells wer processed for RNA isolation and qPCR for the detection of iNOS, IL1β, IL6 and MCP1. The data presented are the mean ± SD of 3 values. *p < 0.05, **p < 0.01, and ***p < 0.001. student’s t test, one-way ANOVA.

Glycolysis has been shown to be an important metabolic pathway that regulates inflammation in myeloid cells. Since, we observed that 2DG treated animals showed altered inflammatory phenotype of CD4 T cell and monocytes/macrophages, we examined the *in vitro* effect of glycolysis on inflammation of mixed glial cells under inflammatory conditions (LPS/IFNγ; LI, treatment). We used various doses of 2DG ranging from 1 to 100 mM and examined the ECAR profile in mixed glial cells and found that lower dose of 2DG reduced the glycolysis in a dose dependent manner (**Figure 7F**). Treatment with 2DG inhibited the production of NO and the expression of iNOS in a dose dependent manner (**Figure 7G**). 2DG treatment also significantly abrogated the messenger RNA expression of iNOS, IL1β (**Figure 7H**) and MCP1 (data not shown), suggesting that inhibition of glycolysis by 2DG results in an anti-inflammatory process. To further support the role of glycolysis, we applied the approach of modulating glucose levels to inhibit glycolysis. We cultured brain primary glial cells in low (1 mM) and normal glucose (10 mM) RPMI in presence of normal glutamine levels (2 mM). Mixed glial cells cultured in low glucose exhibited significantly lower glycolysis compared to the cells cultured with normal glucose as expected (**Figure 7I**). When cells were treated with LPS/IFNγ (LI), the low glucose condition (1 mM) reduced the expression of iNOS by more than 10-fold compared to that observed under normal glucose condition (10 mM) (**Figure 7J**). Low glucose is known to activate the cellular energy sensor, AMP-activated protein kinase (AMPK), which has emerged as a metabolic regulator of energy metabolism and the ensuing functional activity of various immune cells. We observed higher phosphorylation of AMPK and its endogenous substrate, acetyl-CoA carboxylase, in the mixed glial cells cultured in lower glucose compared to glial cells in normal glucose levels (**Figure 7J**). In addition, inhibition of glycolysis by glucose deprivation also resulted in lower expression of iNOS, IL1β, IL6 and MCP1 compared to glial cells with normal glucose upon LI stimulation (**Figure 7K).** Overall, these data suggest that targeting glycolysis either by pharmacological approach (2DG) or by substrate (glucose) deprivation results in reduction in inflammatory response in brain glial cells and may be the underlying mechanism of the protective effect of 2DG in MS/EAE.

## DISCUSSION

MS is a disease where interplay between inflammation and immune cells dictates the disease progression. Recent advances have now established a strong link between the infiltrated immune cell function and altered cellular metabolism (Cohan et al., 2020; Makowski et al., 2020). In the present study, we adapted the unique approach of interrogating the serum metabolite profile of RRMS patients to arrive at a targetable metabolic pathway. By integrating the patient based untargeted metabolomics and functional metabolism, which were validated by extensive preclinical and ex-vivo experimentation, we report that glycolysis is the key pathway regulating the metabotype and consequently the function of immune cells including CD4 and monocytes/macrophages in RRMS/EAE. Modulating the cellular metabolism by inhibition of glycolysis, can curtail the EAE disease process possibly by reverting immune cells towards an anti-inflammatory phenotype of macrophage and resulting in an anti-inflammatory protective environment.

Our untargeted metabolomics approach revealed 60 significantly altered metabolites in patient samples compared to HS, which strongly suggests the presence of a distinct dysregulated metabolic state in RRMS. Analyzing the biological implications of these altered metabolites by various analytical tools identified SIPR2 (intermediate regulators JUN, IFNβ1, and fingolimod) and TGFβ1 (intermediate regulator IRF7) as master regulators associated with the metabolite changes during the disease state. Studies have shown that sphingolipid metabolites such as S1P and ceramides strongly influence many cellular functions under physiological or pathological conditions (Strub et al., 2010; Zhang et al., 1991; Zheng et al., 2006). S1P is a bioactive lipid that mediates signaling and controls various physiological functions through members of the S1PR subfamily except for SIPR2 (Strub et al., 2010). According to the CNA, S1PR2, IFNβ1, and fingolimod have an inhibitory effect on MS. IFNβ1 is a cytokine in the interferon family used to treat RRMS as one of the first-line drugs. Fingolimod (also known as FTY720) is the first Food and Drug Administration approved oral disease-modifying drug for MS (Brinkmann et al., 2010), and its phosphorylated form is considered as a functional antagonist of S1PR1 (Billich et al., 2003), by which it impacts lymphocyte trafficking and stops infiltration into CNS and mitigates the inflammatory process. The other master regulator identified, TGFβ1 cytokine plays a pleiotropic regulatory role and a synergistic role in disease activation in conjunction with transcription factor IRF7 (Ning et al., 2011), which regulates interferon genes (Mirshafiey and Mohsenzadegan, 2009). Thus, the altered metabolites identified in our study are connected with or are a consequence of the aberrant biological processes that drive the MS disease. Metscape analysis revealed a global metabolomic derangement during MS disease resulting in perturbation of several metabolic networks that include glycolysis, TCA, urea, purine, tyrosine, butnoate, phosphatidylinositol phosphate, glycerophospholipids, glycerosphingolipids, pyrimidine, aminosugar, methionine and cysteine. This observation is supported by some of the previous studies in MS patients and preclinical mouse models of EAE (Mangalam et al., 2013; Poisson et al., 2015; Stoessel et al., 2018). MetaboAnalyst analysis revealed the 4 major pathways encompassing the altered metabolites pertained to glycerophospholipid, citrate, sphingolipid, and pyruvate metabolism, where glycolysis is the common upstream pathway feeding and connecting to these pathways, reaffirming the fact that enhanced glycolysis is a peculiar feature of the autoimmune diseases (Kaushik et al., 2019; Radu et al., 2007; Zhang et al., 2019). Apart from fulfilling the cellular energy expenses, glycolysis has a pivotal role in regulating the proper functioning of cells by generating biosynthetic intermediates and precursors required not only for cellular growth and proliferation, but also normal functioning of any cell.

Recent studies have shown that enhanced glycolysis in various immune cells is a hallmark for carrying out successful immune processes like cell activation, proliferation and effector response (Locasale, 2013). We show that the altered glycolysis related serum metabolites are reflective of the immune cell’s energy status as the immune cells from RRMS patients had inherent higher functional glycolysis. This suggests that PBMCs of RRMS patients have an altered metabotype that relies mainly on glycolysis to meet their increased energy needs in the inflammatory environment. To ascertain if glycolysis is a valid target with potential to treat MS, we adopted a pharmacological approach to inhibit glycolysis and examine its effect on the inhibition of disease progression in EAE. Few studies have suggested glycolytic genes as therapeutic targets, including pyruvate kinase M2 (Angiari et al., 2020; Damasceno et al., 2020; Kono et al., 2019), LDHA (Kaushik et al., 2019) and glucose phosphate isomerase (Wu et al., 2020). We selected a general inhibitor of glycolysis, 2-DG, which acts as a glucose analog, phosphorylated by hexokinase to 2DG6P, and cannot be further metabolized, resulting in revoking glycolysis in cells (Wick et al., 1957). We show that 2DG can offer significant protection in the clinical score, which implies a better clinical course and an overall improvement in various models of relapsing-remitting and chronic EAE. 2DG treatment blocked the infiltration of all MN cells, resulting in the protection of EAE. Previosuly, Shi et al. reported that glycolytic activity in CD4 T cells could contribute to the lineage choices between Th17 and Treg cells and affect the EAE disease outcome (Shi et al., 2011). Here, using 2DG as a glycolytic inhibitor,we did not observe any effect on Tregs, which is in contrast to the previous study, and 2DG may play different roles in vivo vs. *in vitro* polarization condtions (Shi et al., 2011). Here we examined the detailed mechanism of 2DG-mediated protection in various models of EAE. Our most striking observation is that 2DG polarizes monocytes/macrophages into an anti-inflammatory phenotype and increases the ratio of anti-inflammatory to pro-inflammatory macrophages in CNS. It is well established that both pro-inflammatory (M1, the classically activated) and anti-inflammatory (M2, also known as alternatively activated macrophage) population is identified in MS (Boven et al., 2006; Raine, 2017; Zhang et al., 2011). Their dual function is well defined in EAE as peripheral depletion of monocytes by clodronate liposomes reduced the severity of EAE (Huitinga et al., 1990) and the administration of anti-inflammatory monocytes (M2) ameliorates EAE (Mikita et al., 2011; Weber et al., 2007). Monocytes isolated from 2DG treated EAE group could restrict the ongoing EAE disease supporting that anti-inflammatory phenotype of monocytes/macrophages are protective in EAE and MS. Further, the therapeutic potential of M2 monocytes in MS has been highlighted by recent studies showing MS approved drugs, glatiramer acetate, laquinimod, and fasudil, suppress EAE by promoting the development of M2 activated monocytes (Liu et al., 2013; Schulze-Topphoff et al.; Weber et al., 2007). We show that the mechanism of 2DG-mediated promotion of anti-inflammatory phenotype is via the inhibition of glycolysis, since monocytes isolated from 2DG treated mice maintained a lower glycolytic function compared to the untreated EAE group. Higher glycolysis in immune cells during EAE (Kaushik et al., 2019; Wu et al., 2020) and other diseases (Liu et al., 2018; Okano et al., 2017; Weyand et al., 2017; Yin et al., 2016) clearly supports that metabolic reprograming in inflammatory cells is an attractive target for immune therapy in autoimmune disorders. The outcome of our clinical and preclinical studies strongly underlines the role of glycolysis in the regulation and functioning of immune cells during the inflammatory state within the body. This makes it entirely plausible to target the metabolic pathways for developing new therapeutics for autoimmune diseases, including MS.

Overall, through our approach, we identified the distinct metabolite separation between normal and relapsing-remitting subjects and identified 4 significant pathways, with glycolysis being the central connector of these pathways being impacted due to disease in patients. On the one hand, the observed metabolite signature in the biofluid of MS patients can be exploited for biomarker development in the future. On the other hand, bioinformatics revealed a common upstream testable metabolic pathway, glycolysis, which we demonstrate can be successfully targeted in mouse models of MS using a glycolysis inhibitor. The 2DG-mediated inhibition of glycolysis strongly suggests the beneficial effect of this strategy on the disease progression in EAE. Moreover, an in-depth study of the altered pathway(s) could provide insight into the disease’s immunopathological mechanisms. In conclusion, our study provides additional evidence to support the existence of metabolic reprogramming in MS along with other autoimmune diseases with the novel observation of an altered metabotype in the patient’s immune cells, defined by a higher glycolysis, which certainly entails the potential to selectively target the glycolytic pathway for future therapeutic interventions.

## Limitations

MS is a chronic disease, and we recognize the limitation of the 2DG being translated for MS treatment due to poor drug-like characteristics including rapid metabolism and short half-life (Hansen et al., 1984). To enhance its drug-like property, new analogs of 2DG are being tested (Pajak et al., 2019). 2DG has been approved by the Indian Defense Research and Development Organization as adjunct therapy in moderate to severe COVID-19 patients. 2DG-treated COVID-19 patients showed a significant improvement symptomatically and became free from oxygen supplementation by 3 days compared to untreated patients. 2DG is also being extensively tested against various cancers in clinical trials (Pajak et al., 2019). We acknowledge that the therapeutic use of 2DG will have to be specifically designed around inflammatory episodes. Nevertheless, this is a proof-of-the-principle study to demonstrate the implication of metabolomics to identify a therapeutic target. In support, glycolytic inhibitors have been shown to be protective in EAE (Angiari et al., 2020; Damasceno et al., 2020; Kaushik et al., 2019; Kono et al., 2019; Wu et al., 2020), further supporting our metabolomics-driven interpretation in an effort to identify a therapeutic target for MS.

## STAR METHODS SECTION

### Patient recruitment and blood sampling

Blood samples collected from 35 unrelated RRMS patients were recruited for the present study from the Biobank facility at the Department of Neurology, Henry Ford Hospital, Detroit, MI. Patients were diagnosed for MS, according to the 2010 revised McDonald criteria by Dr. Cerghet (Polman et al., 2011). The portion of the study on patient samples stands approved by the Institutional Review Board of Henry Ford Health System. All the patients/guardians were informed of the study, and informed consent was acquired from them in accordance with the ethical standards laid down by the World Health Organization and Declaration of Helsinki 1964 and its later amendments or comparable ethical standards. Inclusion criteria for selecting patients included radiological confirmed MS patients, while exclusion criteria included patients who did not satisfy McDonald diagnostic criteria, patients with previous or family history of other neurodegenerative or inflammatory diseases, and patients who rebutted from participating in this study.

Simultaneously, blood samples from 14 age- and gender-matched HS (to confine the confounding effect) were also used to serve as controls. Inclusion criteria for the selection of controls included healthy individuals with minor neurological problems like back pain and headache and those with no previous or family history of MS or any other autoimmune diseases. Exclusion criteria included individuals who denied participating in the study and with previous self and family history of inflammatory disease.

### Serum and PBMCs

For serum analysis, blood samples from RRMS patients and HS were collected in red-capped vacutainer tubes. Serum was separated from blood by centrifugation at 1500 rpm for 10 minutes. The clear yellow liquid supernatant part was collected from the top and stored in respective yellow capped serum tubes at −80°C till further processing. Simultaneously, PBMCs were isolated from blood using the Ficoll-Hypaque gradient method. Blood collected in ethylene diamine tetraacetic acid coated vacutainer tubes were mixed with an equal volume of 1X sterile PBS (in the ratio of 1:1), and it was subsequently transferred onto the equal volume of Ficoll contained in 15-ml falcon tube. After that, the samples were centrifuged at 3000 rpm with zero deceleration for 30 minutes. Ficoll sets up the gradient, and the PBMC layer was separated between the topmost plasma and Ficoll. The collected PBMCs were again washed with 1X PBS at 1500 rpm for 10 minutes, and then cells were counted using BioRad Cell Counter (Hercules, CA). They were stored in liquid nitrogen until further use in 10% fetal bovine serum and 10% DMSO. The details of peptides and reagents are given in Star Methods.

### Animals

Female B6, SJL (10-12 weeks old) and 2D2 TCR (MOG_35-55_ specific TCR transgenic mice (Stock 006912) were purchased from the Jackson Laboratory (Bar Harbor, ME). Animals were housed in the pathogen-free animal facility of Henry Ford Hospital, Detroit, MI, according to the animal protocols approved by the Animal Care and Use Committee of Henry Ford Hospital.

### EAE induction and recall response

B6, 2D2 and SJL mice (10-12 weeks old) were immunized on day 0 by subcutaneous injections in the flank region with a total 200 μl of emulsion containing antigen MOG_35-55_ or PLP_139-151_ peptide (100 μg/mouse), along with killed Mycobacterium tuberculosis H37Ra (400 μg) in CFA as described previously (Mangalam et al., 2013; Mangalam et al., 2016; Poisson et al., 2015). Pertussis toxin at the dose of 300 ng/mouse in PBS was given to B6 or 2D2 immunized mice on day 0 and 2 post-immunization in the volume of 200 μl. Pertussis toxin was not injected in SJL mice. One set of mice were injected with CFA/pertussis toxin without antigen/peptide named as control. Clinical disease was monitored daily in a blinded fashion by measuring paralysis according to the conventional grading system: 0, no disease; 1, complete loss of tail tonicity; 2, partial hind limb paralysis (uneven gate of hind limb); 3, complete hind limb paralysis; 4, complete hind and forelimb paralysis; and 5, moribund or dead. Furthermore, cells (4×10^6^/ml) isolated from spleens were cultured in the presence or absence of antigen (20 μg/ml). Cell proliferation and the production of various cytokines (IFNγ, GM-CSF, and IL17) were examined as described before (Mangalam et al., 2013; Mangalam et al., 2016; Poisson et al., 2015). Cells were also processed for cell surface and intracellular staining.

### Histopathology and Luxol fast blue staining

Hematoxylin and eosin and Luxol fast blue staining methods and histopathological analyses protocols were adopted from our reports (Khan et al., 2015; Mangalam et al., 2016; Vaibhav et al., 2018). Mice were anesthetized with isoflurane and transcardially perfused with 0.9% chilled 1X PBS followed by 4% paraformaldehyde. Mouse spinal cord tissues were harvested and fixed in 4% paraformaldehyde for 24 hours at 4°C and later embedded in paraffin. Briefly, 5-µm spinal cord sections were obtained and stained with hematoxylin and eosin to check the infiltration of immune cells and the analysis of lesion size. Loss in white matter content and subsequent demyelination was visualized with Luxol fast blue staining. Brightfield images were captured using a light microscope, the demyelinated area was selected, measured and expressed as percentage of total area ([demyelinated area/total spinal cord area] × 100). Immunohistochemistry was performed on paraffin embedded spinal cord sections. Briefly, sections were deparaffinized, blocked and incubated with specific primary antibody (to CD4 and F4/80) overnight at 4°C. Sections were then washed 3 times with 0.1% PBST followed by incubation with an appropriate secondary antibody for 2 hours at room temperature. Thereafter, sections were rinsed, incubated with ABC reagent for 2 hours and visualized with 3,3′-diaminobenzidine. Images were acquired using a bright field microscope, converted into 8-bit images, threshold filtered and quantified using ImageJ software (National Institutes of Health, Bethesda, MD).

### Flow cytometry

Surface markers of CNS-infiltrating or spleen cells were stained by incubating the cells with fluorochrome-labeled antibodies against Ly6C, F4/80, CD38 and EGR2 for 30 minutes at 4°C. MOG_35-55_-specific activation of Th1 and Th17 from CNS-infiltrating lymphocytes were analyzed by stimulating the cells with the MOG_35-55_ peptide (20 ug/ml) for 18 hours followed by treatment with GolgiPlug at 37°C. After 5-hours of incubation, cells were harvested, washed, and stained with fluorochrome-labeled antibodies against cell surface markers CD3, CD4 and CD45. Subsequently, for intracellular markers staining, cells were fixed and permeabilized using cytofix/cytoperm buffer (Biolegend, San Diego, CA) and permeabilization buffer (Biolegend) followed by incubation with monoclonal antibodies against IFNγ and IL17 and GM-CSF for 30 minutes at room temperature. Data for flow cytometry were acquired on BD LSR II (BD Biosciences, Franklin Lakes, NJ), and results were analyzed using FACSDiva software (BD Biosciences, Franklin Lakes, NJ).

### Quantitative PCR (qPCR) and NCBI GEO database analysis

Total RNA was isolated from isolated monocytes of treated and untreated groups with 2DG using Direct-zol RNA kit (Zymo Research, Irvine, CA). The complementary DNA was prepared using iScript cDNA synthesis kit (Bio-Rad) as per the manufacturer’s protocol. qPCR was performed using SYBER Green PCR master mix (Bio-Rad) and a Bio-Rad iCycler iQ PCR (Bio-Rad) using specific primers against specific genes including iNOS, IL1β, arginase 1, chitinase-like 3 (Ym1/2), glucose transporter 1, hexokinase 2, triose-phosphate isomerase, pyruvate kinase M, LDHA and monocarboxylate transporter 1. Expression of glycolytic genes were normalized with the control gene, ribosomal L27 housekeeping gene.We selected 3 gene expression profile datasets about MS, GSE21942 (Kemppinen et al., 2011), GSE26484 (Nakatsuji et al., 2012), and GSE43591 (Jernas et al., 2013). All 3 datasets are based on Affymetrix. Raw data of each dataset is downloaded from NCBI GEO database with GEOquery in R and is preprocessed with RMA algorithm. We performed differential expression analysis with Limma between the MS group and the HS on each dataset separately.

### Glucose, lactate, and ATP measurements

The glucose and lactate levels in media were measured using Glucose Glo and Lactate Glo assay kits (Promega) as per the manufacturer’s instructions. In brief, isolated monocytes using EasySep mouse monocyte isolation kit (Stemcell Technologies, Cambridge, MA) from the spleen of treated and untreated EAE with 2DG were plated (25K/well) in a 96-well plate in RPMI media containing 10 mM glucose, 2 mM glutamine and 5% dialyzed fetal bovine serum. After 24 hours, culture media was processed for the measurement of glucose and lactate using their respective assay kits from Promega. Total cellular levels of ATP were measured using ATPlite luminescence assay kit from Perkin Elmer. In brief, isolated monocytes (10,000 cells per well) were plated in a 96-well plate in 50 ul of PBS. Cells were lysed by adding 50 ul of mammalian cell lysis buffer, and the plate was shaken for 5 minutes at room temperature. Fifty microliters of substrate were added, and the plate was shaken in the dark for 5-10 minutes before measuring the luminescence. Ten millimolar of the ATP solution was used to set up an ATP standard curve, and all samples were within the linear range of the assay.

### Extracellular Acidification Rate (ECAR)

Seahorse Bioscience XF^e^96 Extracellular Flux Analyzer (Agilent) was used to measure the glycolytic activity of intact PBMCs by monitoring the ECAR in media. ECAR was measured as described before (Dar et al., 2017). In brief, the XFe96 sensor cartridge was soaked in Calibrant overnight. PBMCs from HS and RRMS patients or CFA and EAE groups were seeded with a concentration of 0.1×10^6^ cells per well in poly-lysine coated XF^e^ 96-well seahorse culture microplate in 100µl of Agilent Seahorse XF DMEM medium, pH 7.4 containing 2mM glutamax (Thermofisher). The plate was centrifuged for 60 seconds at 400 RPMI. Seventy-five microliter of DMEM media was added in seahorse culture microplate and kept at a CO_2_-free incubator at 37°C for degassing. Injection of glucose as fuel via first port induced the ECAR levels in cells. The basal glycolytic activity was calculated as the difference between ECAR reading following the pre-injection and the injection of glucose. Blanks without cells were automatically subtracted during analysis by Agilent seahorse XF technology. ECAR values were normalized to cell number.

### Metabolomic analysis

Metabolomic profiling analysis was performed by Metabolon Inc. (Durham, NC), as previously described (Mangalam et al., 2013; Poisson et al., 2015; Singh et al., 2019).

#### Sample accessioning

Each sample received was accessioned into the Metabolon LIMS system and was assigned by the LIMS, a unique identifier that was associated with the original source identifier only. This identifier was used to track all sample handling, tasks, and results. The samples (and all derived aliquots) were tracked by the LIMS system. All portions of any sample were automatically assigned their own unique identifiers by the LIMS when a new task was created; these samples’ relationship was also tracked. All samples were maintained at −80°C until processed.

#### Sample preparation

Samples were prepared using the automated MicroLab STAR system from Hamilton Company. A recovery standard was added prior to the first step in the extraction process for QC purposes. To remove protein, dissociate small molecules bound to protein or trapped in the precipitated protein matrix, and to recover chemically diverse metabolites, proteins were precipitated with methanol under vigorous shaking for 2 minutes (Glen Mills GenoGrinder 2000) followed by centrifugation. The resulting extract was divided into 5 fractions: one for analysis by ultra-performance liquid chromatography-mass spectroscopy (UPLC-MS/MS) with positive ion mode electrospray ionization, one for analysis by UPLC-MS/MS with negative ion mode electrospray ionization, one for analysis by UPLC-MS/MS polar platform (negative ionization), one for analysis by gas chromatography-mass spectroscopy (GC-MS), and one sample was reserved for backup. Samples were placed briefly on a TurboVap (Zymark) to remove the organic solvent. For LC, the samples were stored overnight under nitrogen before preparation for analysis. For GC, each sample was dried under vacuum overnight before preparation for analysis.

#### Ultrahigh performance liquid chromatography-tandem mass spectroscopy (UPLC-MS/MS)

The LC/MS portion of the platform was based on a Waters ACQUITY UPLC, and a Thermo Scientific Q-Exactive high resolution/accurate mass spectrometer interfaced with a heated electrospray ionization (HESI-II) source and Orbitrap mass analyzer operated at 35,000 mass resolution. The sample extract was dried then reconstituted in acidic or basic LC-compatible solvents, each of which contained 8 or more injection standards at fixed concentrations to ensure injection and chromatographic consistency. One aliquot was analyzed using acidic positive ion optimized conditions and the other using basic negative ion optimized conditions in 2 independent injections using separate dedicated columns (Waters UPLC BEH C18-2.1×100 mm, 1.7 µm). Extracts reconstituted in acidic conditions were gradient eluted from a C18 column using water and methanol containing 0.1% formic acid. The basic extracts were similarly eluted from C18 using methanol and water, however, with 6.5mM Ammonium Bicarbonate. The third aliquot was analyzed via negative ionization following elution from a HILIC column (Waters UPLC BEH Amide 2.1×150 mm, 1.7 µm) using a gradient consisting of water and acetonitrile with 10mM Ammonium Formate. The MS analysis alternated between MS and data-dependent MS^2^ scans using dynamic exclusion, and the scan range was from 80-1000 *m/z*. Raw data files are archived and extracted as described below.

#### Gas chromatography-mass spectroscopy (GC-MS)

The samples destined for analysis by GC-MS were dried under vacuum for a minimum of 18 h prior to being derivatized under dried nitrogen using bistrimethyl-silyltrifluoroacetamide. Derivatized samples were separated on a 5% diphenyl/95% dimethyl polysiloxane fused silica column (20 m x 0.18 mm ID; 0.18 um film thickness) with helium as a carrier gas and a temperature ramp from 60° to 340°C in a 17.5 min period. Samples were analyzed on a Thermo-Finnigan Trace DSQ fast-scanning single-quadrupole mass spectrometer using electron impact ionization (EI) and operated at unit mass resolving power. The scan range was from 50–750 m/z. Raw data files are archived and extracted as described below.

#### Quality assurance/quality control

Several types of controls were analyzed in concert with the experimental samples: a pooled matrix sample generated by taking a small volume of each experimental sample (or alternatively, use of a pool of well-characterized human plasma) served as a technical replicate throughout the data set; extracted water samples served as process blanks; and a cocktail of QC standards that were carefully chosen not to interfere with the measurement of endogenous compounds were spiked into every analyzed sample, allowed instrument performance monitoring and aided chromatographic alignment. Instrument variability was determined by calculating the median relative standard deviation (RSD) for the standards that were added to each sample prior to injection into the mass spectrometers. Overall process variability was determined by calculating the median RSD for all endogenous metabolites (i.e., non-instrument standards) present in 100% of the pooled matrix samples. Experimental samples were randomized across the platform run with QC samples spaced evenly among the injections.

#### Data extraction and compound identification

Raw data was extracted, peak-identified, and QC processed using Metabolon’s hardware and software. These systems are built on a web-service platform utilizing Microsoft’s.NET technologies, which run on high-performance application servers and fiber-channel storage arrays in clusters to provide active failover and load-balancing. Compounds were identified by comparison to library entries of purified standards or recurrent unknown entities. Metabolon maintains a library based on authenticated standards that contain the retention time/index (RI), mass to charge ratio (*m/z)*, and chromatographic data (including MS/MS spectral data) on all molecules present in the library. Furthermore, biochemical identifications are based on 3 criteria: retention index within a narrow RI window of the proposed identification, accurate mass match to the library +/-0.005 amu, and the MS/MS forward and reverse scores between the experimental data and authentic standards. The MS/MS scores are based on a comparison of the ions present in the experimental spectrum to the ions present in the library spectrum. While there may be similarities between these molecules based on one of these factors, the use of all 3 data points can be utilized to distinguish and differentiate biochemicals. More than 3300 commercially available purified standard compounds have been acquired and registered into LIMS for distribution to both the LC-MS and GC-MS platforms for the determination of their analytical characteristics. Additional mass spectral entries have been created for structurally unnamed biochemicals, which have been identified by virtue of their recurrent nature (both chromatographic and mass spectral). These compounds have the potential to be identified by the future acquisition of a matching purified standard or by classical structural analysis.

#### Curation

A variety of curation procedures were carried out to ensure that a high-quality data set was made available for statistical analysis and data interpretation. The QC and curation processes were designed to ensure accurate and consistent identification of true chemical entities and to remove those representing system artifacts, misassignments, and background noise. Metabolon data analysts use proprietary visualization and interpretation software to confirm the consistency of peak identification among the various samples. Library matches for each compound were checked for each sample and corrected if necessary.

#### Metabolite quantification and data normalization

Peaks were quantified using area-under-the-curve. For studies spanning multiple days, a data normalization step was performed to correct variation resulting from instrument inter-day tuning differences. Essentially, each compound was corrected in run-day blocks by registering the medians to equal one (1.00) and normalizing each data point proportionately (termed the “block correction”). For studies that did not require more than one day of analysis, no normalization is necessary, other than for purposes of data visualization. In certain instances, biochemical data may have been normalized to an additional factor (e.g., cell counts, total protein as determined by Bradford assay, osmolality, etc.) to account for differences in metabolite levels due to differences in the amount of material present in each sample.

### Statistical analysis

Metabolite intensities (scaled imputed) obtained from Metabolon. Statistical analyses are (Chong et al., 2019). The first 2 components of a principal components analysis (PCA) were plotted to identify whether samples or batch effects have contributed disproportionally to the variance in the data. Partial least-squares discriminant analysis was used for the assessment of the separability of the samples by supervised clustering of samples based on metabolites intensities. Per metabolite, comparisons were made using two-sample t-tests on the study cohort. Significant differences were determined at p < 0.05. False discovery rates were calculated by the Q value method from the Bioconductor R package and are provided for reference. Heatmaps are drawn for significantly differential metabolites using blue (low) to yellow (high) coloring to depict standardized intensity differences from the metabolite-level mean. Hierarchical clustering on Pearson’s correlation coefficient is used to generate all dendrograms.

Metabolites were mapped into the IPA software using HMDB IDs and KEGG knowledgebase of compound IDs, PubChem, CAS IDs, and networks of altered metabolites were constructed using Fisher exact test including CNA (Kramer et al., 2014). The goal of IPA’s CNA is to prioritize regulatory networks that explain observed biological changes based on altered metabolites. Its algorithm constructs a multi-edge path from a regulator molecule to regulated metabolites. To construct a gene-enzyme-compound network, Metscape-based on the KEGG database was performed to build a metabolic network on significantly altered metabolites. In Metscape, the construction of the metabolic Network pathway is based on HUMDB, KEGG, and EHMN databases (Karnovsky et al., 2012).

## ACKNOWLEDGMENTS

This work is in-part supported by research grants from the National Multiple Sclerosis Society (US) (RG-1807-31964, RG-1508-05912), the US National Institutes of Health (NS112727, AI144004) and Henry Ford Hospital Internal support (A10270, A30967) to SG. AK is supported by R01EY026964. The funders had no role in study design, data collection, and interpretation, or the decision to submit the work for publication.

## AUTHOR CONTRIBUTIONS

IZ wrote the manuscript, HS performed EAE experiments, LMP and ID performed bioinformatics analysis, JW performed immunoblot and finalized the manuscript, MEA and MNH analyzed IHC data and finalized the manuscript, MC, LMP, AK, RR, AKM, contributed the design and execution of experiments and finalized the manuscript. SG conceive the idea, directed the study and designed experiments and finalized the manuscript.

## DECLARATION OF INTERESTS

The authors declare no competing interests.

## Notes

### Competing Interest Statement

The authors have declared no competing interest.

